## Supplemental Material for "BashTheBug: a crowd of volunteers reproducibly and accurately measure the minimum inhibitory concentrations of 13 antitubercular drugs from photographs of 96-well broth microdilution plates"

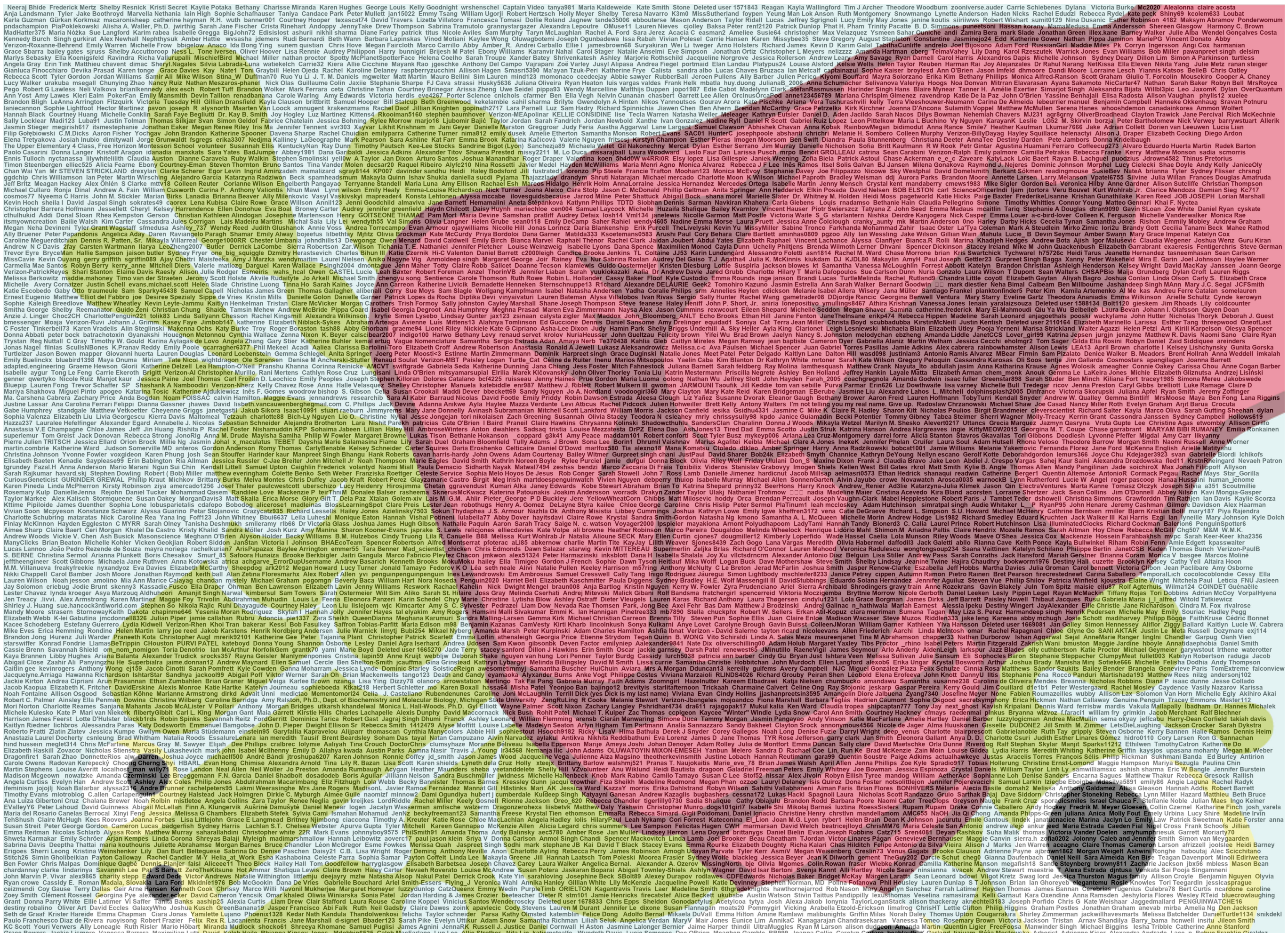

Figure S1: Thank you to all the volunteers who contributed one or more classifications to this manuscript. There are the 5,810 usernames of all the volunteers in this montage – volunteers who did not register or sign in are not included.

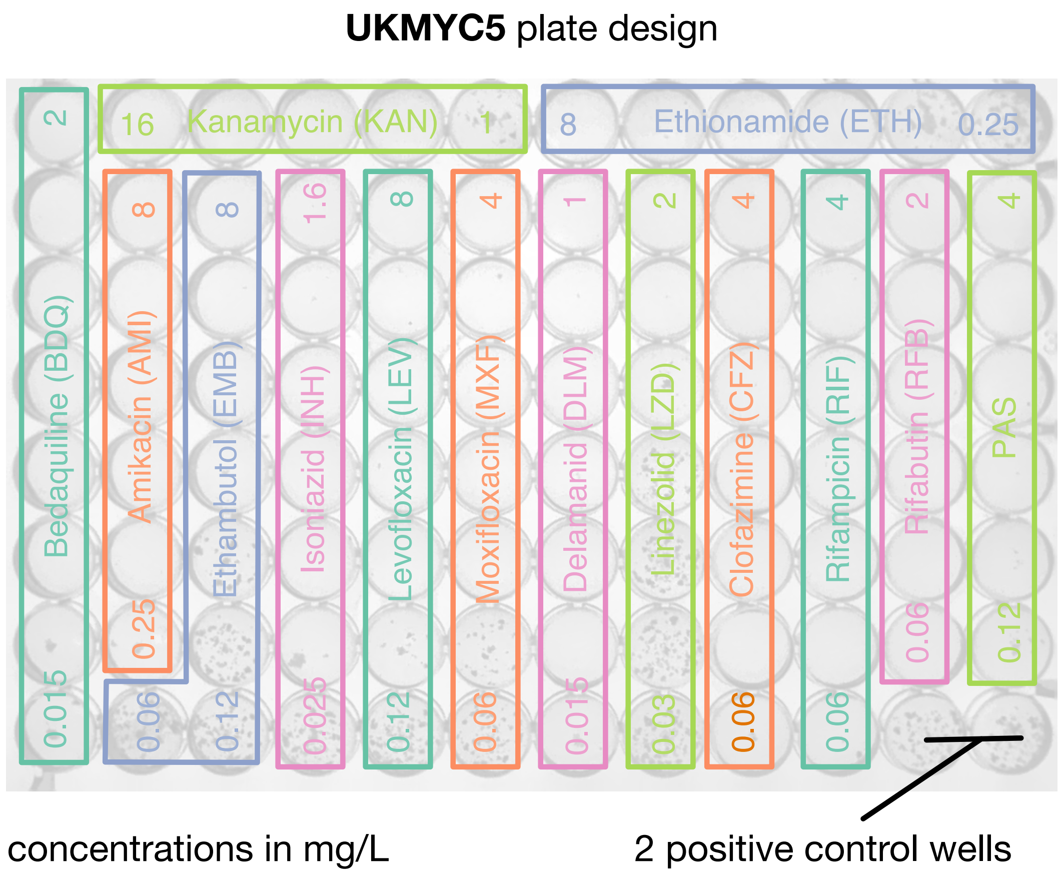

Figure S2: The UKMYC5 plate contains 14 different anti-TB drugs. A previous study[2] showed that *para*-aminosalicylic acid (PAS) performed poorly and it has been removed from the subsequent UKMYC6 plate design. We have therefore excluded this drug from all analyses. Each drug was contained in 5, 6, 7 or 8 wells with each well having double the concentration of drug as the one before. The concentration of the first and last well in each drug series is labelled (mg/L). Two wells contain no drug and are therefore positive control wells.

Table S1: List of strains and their repeats tested by laboratory.

| EQA strain | Vial | Lab Replicate | A | B | C | D | E | G | H |
|---|---|---|---|---|---|---|---|---|---|
| WHO-1 | CRY-19 | 0001 | Y | Y | Y | Y | Y | Y | Y |
| | | 0002 | Y | Y | Y | Y | Y | Y | Y |
| | CRY-7 | 0001 | Y | Y | Y | Y | - | Y | Y |
| | | 0002 | Y | Y | Y | Y | - | Y | Y |
| WHO-10 | CRY-18 | 0001 | Y | Y | Y | - | - | Y | Y |
| | | 0002 | Y | Y | Y | - | - | Y | Y |
| WHO-11 | CRY-16 | 0001 | Y | Y | Y | Y | Y | Y | Y |
| | | 0002 | Y | Y | Y | Y | Y | Y | Y |
| WHO-12 | CRY-13 | 0001 | Y | Y | Y | - | Y | Y | Y |
| | | 0002 | Y | Y | Y | - | Y | Y | Y |
| | CRY-8 | 0001 | Y | Y | Y | Y | Y | Y | Y |
| | | 0002 | Y | Y | Y | Y | Y | Y | Y |
| WHO-13 | CRY-10 | 0001 | Y | Y | Y | - | Y | Y | Y |
| | | 0002 | Y | Y | Y | - | Y | Y | Y |
| | CRY-5 | 0001 | Y | - | Y | Y | Y | Y | Y |
| | | 0002 | Y | - | Y | Y | Y | Y | Y |
| WHO-14 | CRY-29 | 0001 | Y | Y | Y | - | Y | Y | Y |
| | | 0002 | Y | Y | Y | - | Y | Y | Y |
| | CRY-6 | 0001 | Y | Y | Y | Y | Y | Y | Y |
| | | 0002 | Y | Y | Y | Y | Y | Y | Y |
| WHO-15 | CRY-2 | 0001 | Y | Y | Y | Y | Y | Y | Y |
| | | 0002 | Y | Y | Y | Y | Y | Y | Y |
| | CRY-25 | 0001 | Y | Y | Y | - | Y | Y | Y |
| | | 0002 | Y | Y | Y | - | Y | Y | Y |
| WHO-16 | CRY-12 | 0001 | Y | Y | Y | Y | Y | Y | Y |
| | | 0002 | Y | Y | Y | Y | Y | Y | Y |
| | CRY-24 | 0001 | Y | Y | Y | - | Y | Y | Y |
| | | 0002 | Y | Y | Y | - | Y | Y | Y |
| WHO-17 | CRY-14 | 0001 | Y | Y | Y | Y | - | Y | Y |
| | | 0002 | Y | Y | Y | Y | - | Y | Y |
| WHO-18 | CRY-20 | 0001 | Y | Y | Y | Y | Y | Y | Y |
| | | 0002 | Y | Y | Y | Y | Y | Y | Y |
| WHO-19 | CRY-23 | 0001 | Y | Y | Y | Y | Y | Y | Y |
| | | 0002 | Y | Y | Y | Y | - | - | Y |

Continued on next page

Table S1: List of strains and their repeats tested by laboratory.

| EQA strain | Vial | Lab / Replicate | A | B | C | D | E | G | H |
|---|---|---|---|---|---|---|---|---|---|
| WHO-2 | CRY-26 | 0001 | Y | Y | Y | Y | Y | Y | Y |
| | | 0002 | Y | Y | Y | Y | Y | Y | Y |
| WHO-3 | CRY-30 | 0001 | Y | Y | Y | - | Y | Y | Y |
| | | 0002 | Y | Y | Y | - | Y | Y | Y |
| | CRY-4 | 0001 | Y | Y | Y | Y | Y | Y | Y |
| | | 0002 | Y | Y | Y | Y | Y | Y | Y |
| WHO-4 | CRY-22 | 0001 | Y | - | Y | - | Y | Y | Y |
| | | 0002 | Y | - | Y | - | Y | Y | Y |
| | CRY-9 | 0001 | Y | - | Y | Y | Y | Y | Y |
| | | 0002 | Y | - | Y | Y | Y | Y | Y |
| WHO-5 | CRY-15 | 0001 | - | Y | Y | Y | Y | Y | Y |
| | | 0002 | - | Y | Y | Y | Y | Y | Y |
| | CRY-21 | 0001 | Y | Y | Y | - | Y | - | Y |
| | | 0002 | Y | Y | Y | - | Y | Y | Y |
| WHO-6 | CRY-11 | 0001 | Y | Y | Y | - | Y | Y | Y |
| | | 0002 | Y | Y | Y | - | Y | Y | Y |
| | CRY-3 | 0001 | Y | Y | Y | - | Y | Y | Y |
| | | 0002 | Y | Y | Y | - | Y | Y | Y |
| | H37rV | 0001 | Y | Y | Y | Y | Y | Y | Y |
| | | 0002 | Y | Y | Y | Y | Y | Y | Y |
| | | 0003 | Y | Y | Y | Y | Y | Y | Y |
| | | 0004 | Y | Y | Y | Y | Y | Y | Y |
| | | 0005 | Y | Y | Y | Y | Y | Y | Y |
| | | 0006 | Y | Y | Y | Y | Y | Y | Y |
| | | 0007 | Y | Y | Y | Y | Y | Y | Y |
| | | 0008 | Y | Y | Y | Y | Y | Y | Y |
| | | 0009 | Y | Y | Y | Y | Y | Y | Y |
| | | 0010 | Y | Y | Y | Y | Y | Y | Y |
| WHO-7 | CRY-1 | 0001 | Y | Y | Y | Y | - | Y | Y |
| | | 0002 | Y | Y | Y | Y | - | Y | Y |
| | CRY-17 | 0001 | Y | Y | Y | - | Y | Y | Y |
| | | 0002 | Y | Y | Y | - | Y | Y | Y |
| WHO-8 | CRY-27 | 0001 | Y | Y | Y | Y | Y | Y | Y |
| | | 0002 | Y | Y | Y | Y | Y | Y | Y |

Continued on next page

Table S1: List of strains and their repeats tested by laboratory.

| | | Lab | A | B | C | D | E | G | H |
|---|---|---|---|---|---|---|---|---|---|
| EQA strain | Vial | Replicate | | | | | | | |
| WHO-9 | CRY-28 | 0001 | Y | Y | Y | Y | Y | Y | Y |
| | | 0002 | Y | Y | Y | Y | Y | Y | Y |

| Classifications performed by volunteers | Number of drug images | Proportion of drug images |
|---|---|---|
| n $\leq$ 14 | 0 | 0.0 |
| 15 | 26 | 0.1 |
| 16 | 3131 | 7.2 |
| 17 | 34943 | 80.0 |
| 18 | 1863 | 4.3 |
| 19 | 941 | 2.2 |
| 20 | 614 | 1.4 |
| 21 or 22 | 754 | 1.7 |
| 23, 24 or 25 | 730 | 1.7 |
| 25 < n < 29 | 408 | 0.9 |
| 30 < n < 39 | 106 | 0.2 |
| 40 < n < 49 | 22 | 0.1 |
| 50 < n < 74 | 20 | 0.0 |
| 75 < n < 99 | 30 | 0.1 |
| 100 < n < 149 | 36 | 0.1 |
| 150 < n < 199 | 28 | 0.1 |
| 200 < n < 299 | 11 | 0.0 |
| 300 < n < 499 | 8 | 0.0 |

Table S2: The number of classifications performed for each drug image. The retirement limit on the Zooniverse platform was set to 17, however, a subset of images recieved many more classifications.

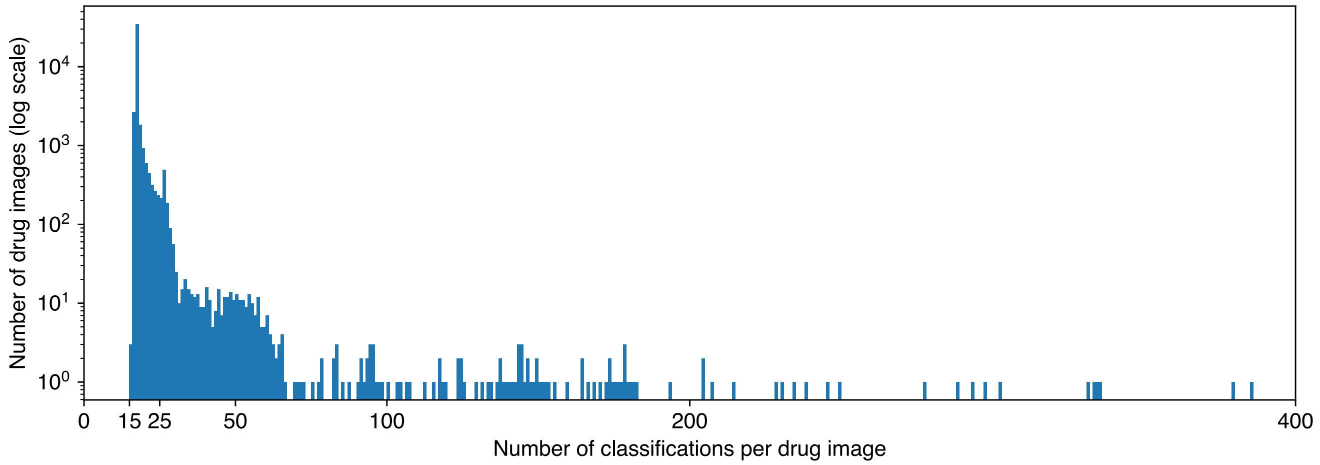

Figure S3: Although the retirement limit within the Zooniverse platform was set to 17, over 1,800 images received more classifications than this and a small number were only classified 15 or 16 times.

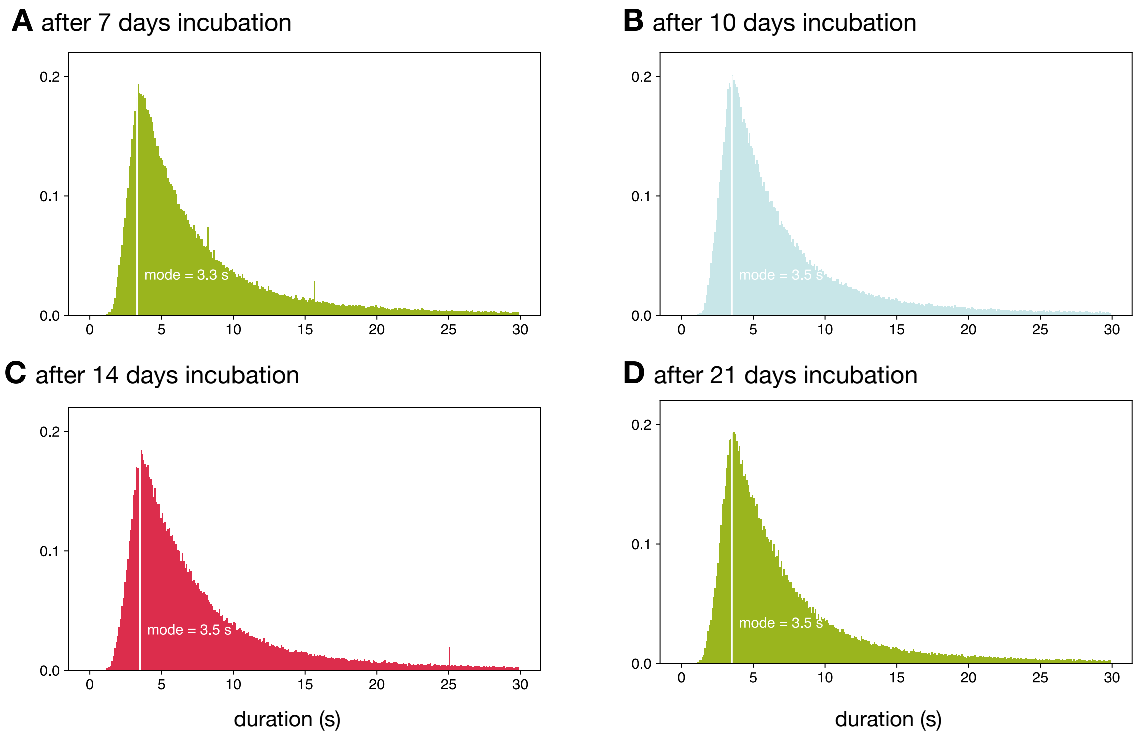

Figure S4: The time spent by volunteers on each classification varied with a mode of 3.5 seconds. Since one would expect different amounts of bacterial growth on the microdilution plates after (**A**) 7, (**B**) 10, (**C**) 14 and (**D**) 21 days the distributions of these were examined separately. All were, however, similar indicating that this did not have a significant effect.

| Reading day | Reference dataset | Measurements | Classifications | Essential agreement | Exact agreement |
|---|---|---|---|---|---|
| 7 | Expert+AMyGDA | 5598 | 80197 | 85.2 % | 68.7 % |
| | Expert | 12502 | 160102 | 81.7 % | 59.9 % |
| 10 | Expert+AMyGDA | 5662 | 85912 | 86.1 % | 73.1 % |
| | Expert | 12474 | 177315 | 82.7 % | 63.2 % |
| 14 | Expert+AMyGDA | 6205 | 112163 | 86.4 % | 74.6 % |
| | Expert | 12488 | 206353 | 83.3 % | 65.3 % |
| 21 | Expert+AMyGDA | 6394 | 106144 | 88.2 % | 78.9 % |
| | Expert | 12474 | 186624 | 85.8 % | 71.1 % |

Table S3: Individual volunteers only agree with the Expert+AMyGDA reference dataset in 60-70% of drug images. The exact and essential agreement between individual volunteers and the reference Expert+AMyGDA dataset improves with the length of incubation. The Expert dataset is shown for comparision.

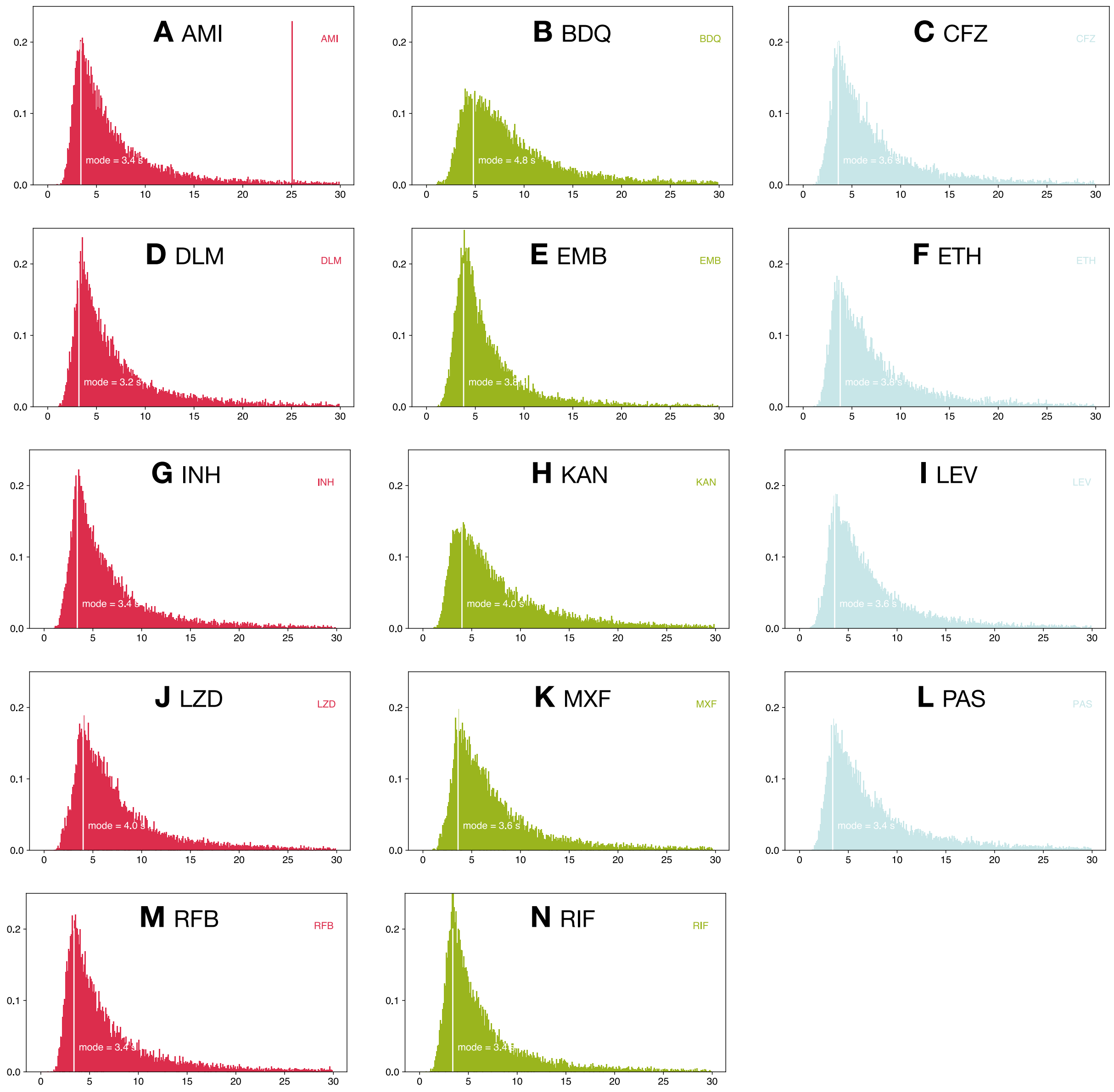

Figure S5: The time spent by volunteers on each classification varied depending on the drug being considered. The drug the volunteers spent the longest on (bedaquiline, mode 4.8 s) was also one of those with the largest number (8) of wells. As measured by its mode of 3.2 s, the volunteers spent the least time classifying delamanid.

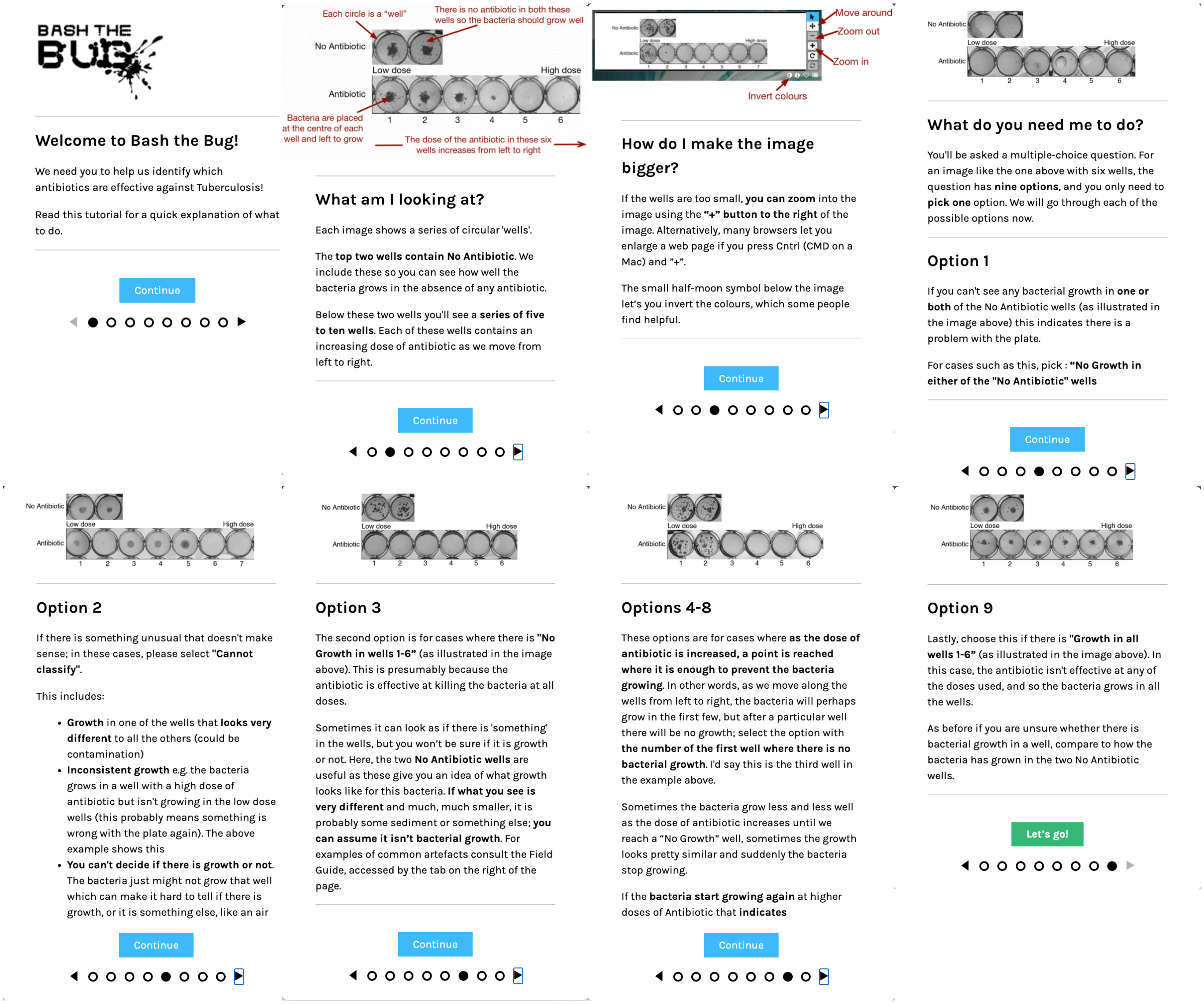

Figure S6: Every new user is shown this tutorial when they first join the BashTheBug Zooniverse project. It uses example images to explain the task and then each of the options that they can choose to classify a drug image.

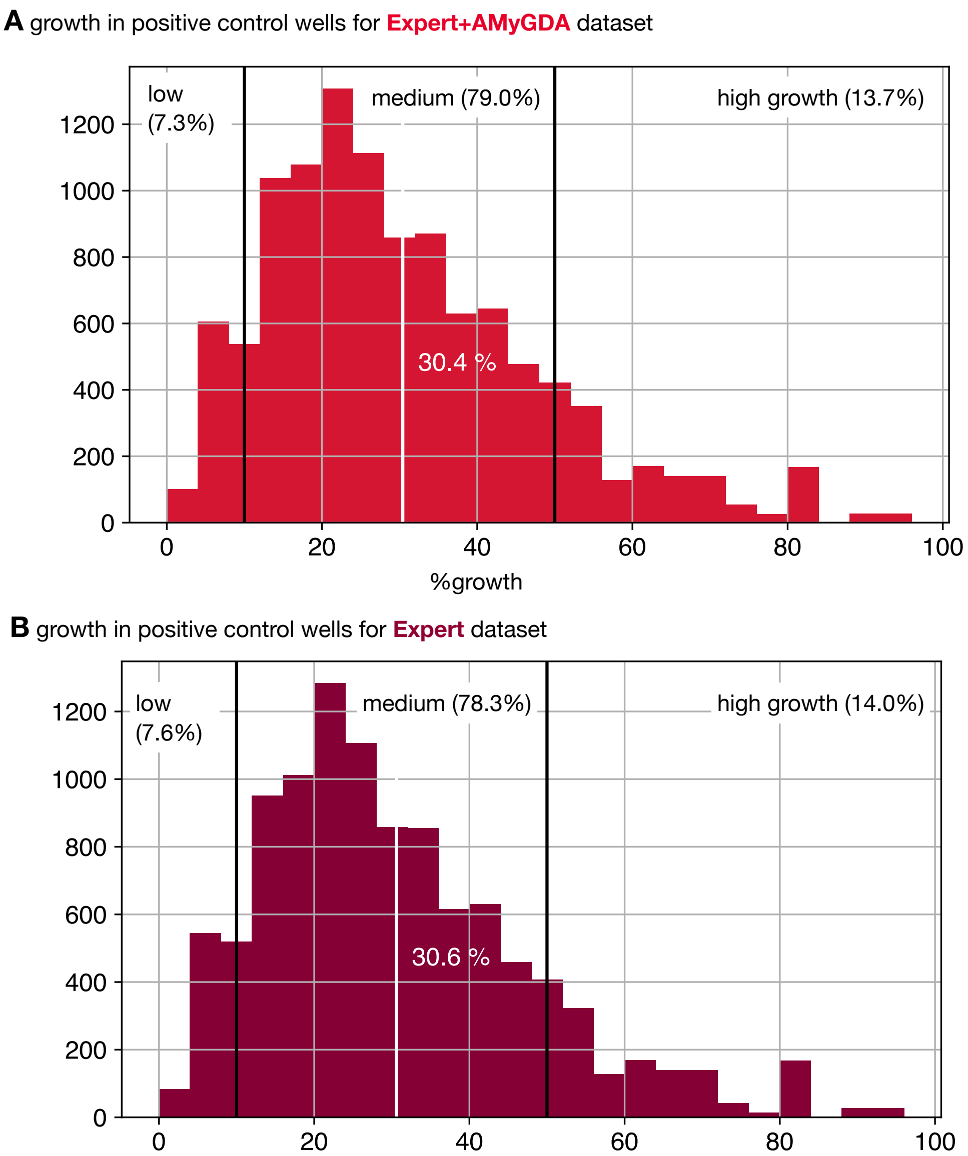

Figure S7: The Expert+AMyGDA consensus dataset has the same distribution of bacterial growth in the positive control wells as the Expert dataset after 14 days incubation. (**A**) The distribution of the mean positive control well growth, as measured by AMyGDA, for the Expert+AMyGDA dataset. The dataset is arbitrarily split into three categories: low ($< 10\%$), medium ($10 \leq$ growth $< 50$ %) and high ($\geq 50$ %) growth. The proportions of the dataset in each category are labelled. (**B**) The distribution of the mean positive control well growth, as measured by AMyGDA, for the Expert dataset. There are around twice as many plates in this dataset (Table S3).

| Dilution | Agreement |
|---:|---|
| NR | 43.9 ± 0.6 % |
| 1 | 76.3 ± 0.4 % |
| 2 | 43.5 ± 0.5 % |
| 3 | 29.1 ± 0.6 % |
| 4 | 31.4 ± 0.8 % |
| 5 | 29.9 ± 0.8 % |
| 6 | 31.6 ± 0.8 % |
| 7 | 39.3 ± 1.0 % |
| 8 | 52.6 ± 1.0 % |
| 9 | 16.1 ± 2.6 % |

Table S4: The Expert and AMyGDA MICs are more likely to concur at smaller dilutions.

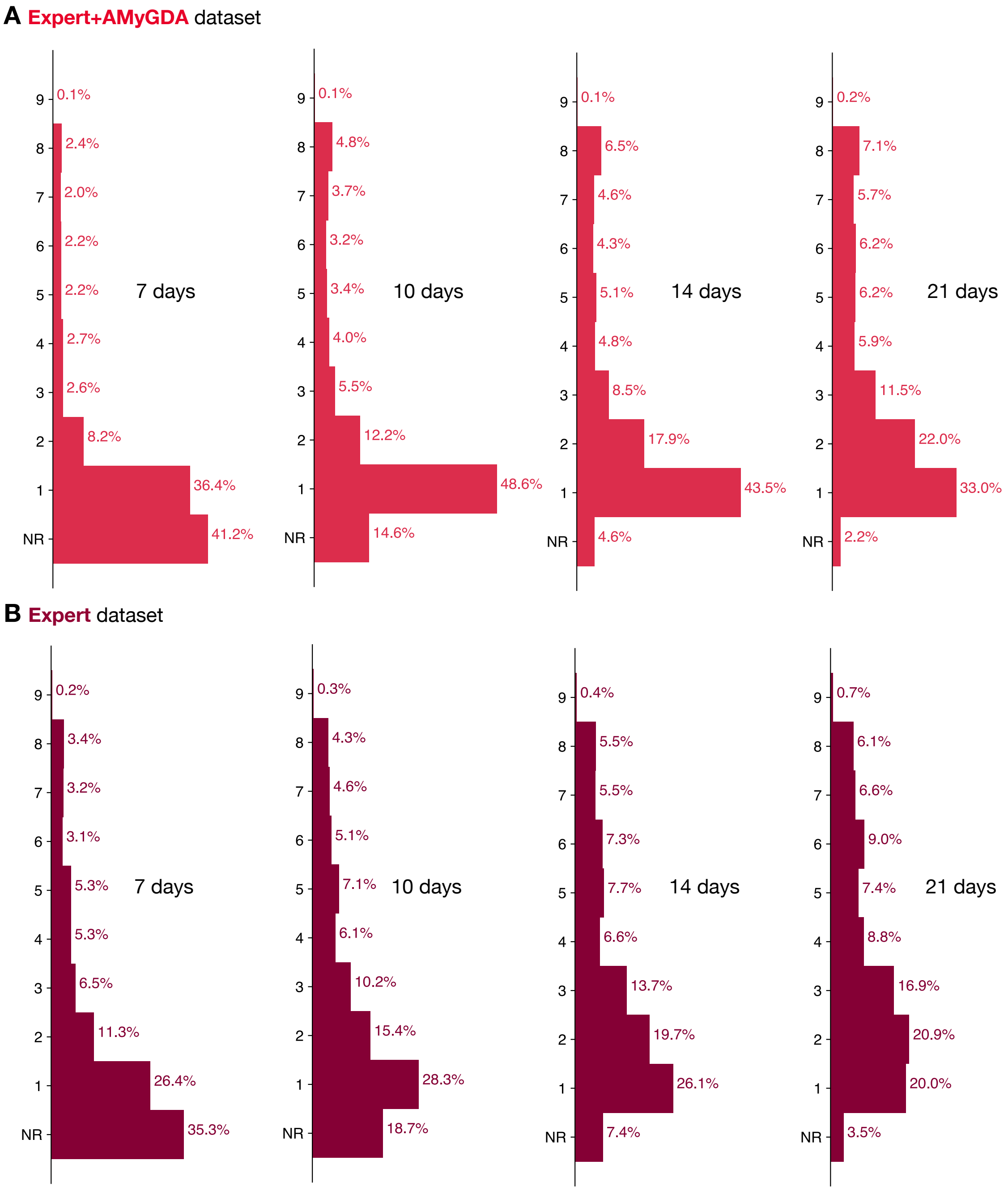

Figure S8: The Expert+AMyGDA dataset has a greater proportion of drug images with low dilutions compared to the Expert dataset. The growth of the bacteria is also evident as the number of days the sample was incubated for is increased.

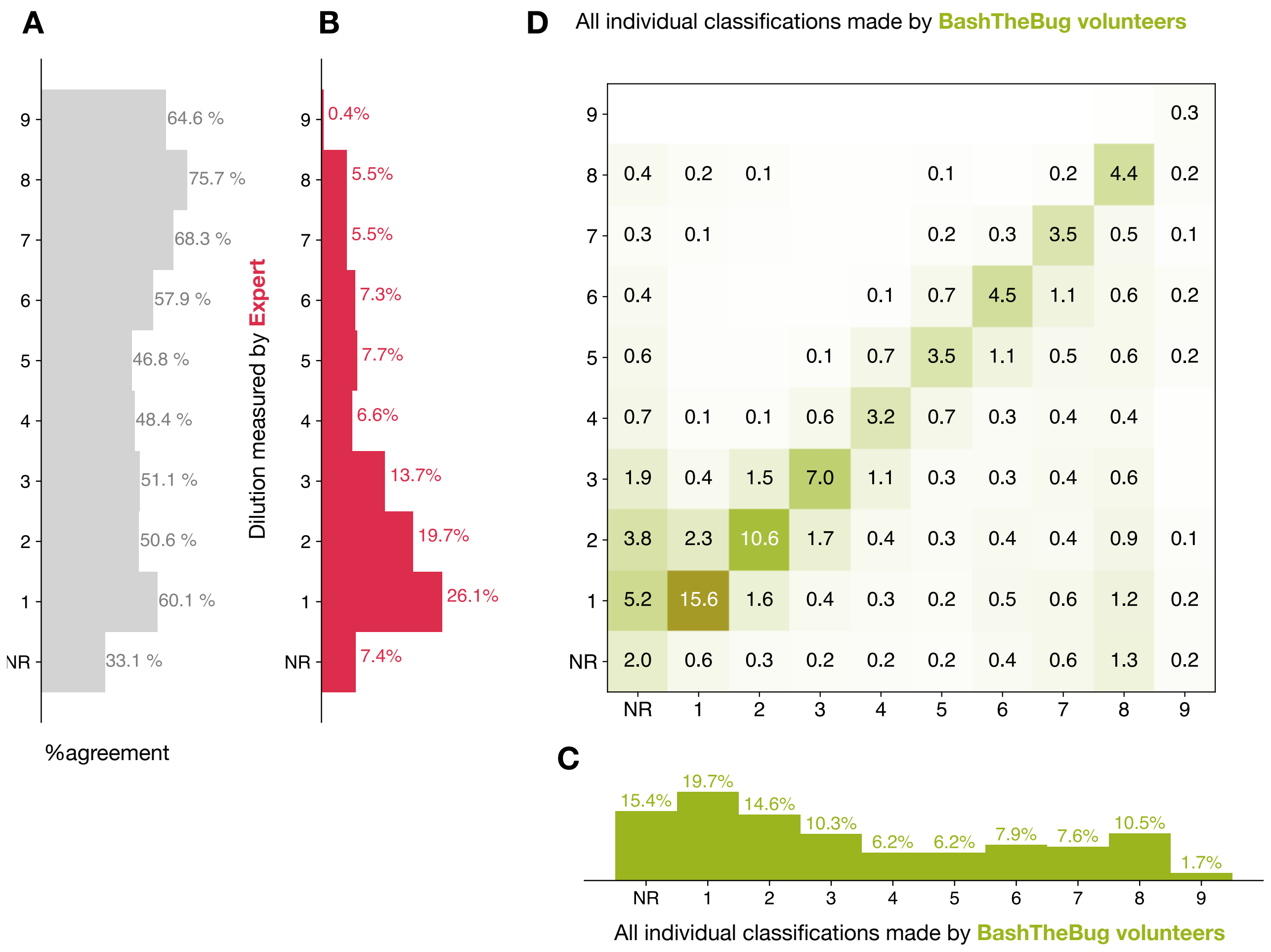

Figure S9: Heatmap showing how all the individual BashTheBug classifications (n=214,164) compare to the set of dilutions where the measurement made by the laboratory scientist using the Thermo Fisher Vizion instrument and a mirrored box after 14 days incubation concur (n=9,402) (**A**) The probability that a single volunteer exactly agrees with the Expert dataset varies with the dilution. The distribution of all MIC dilutions after 14 days incubation read by (**B**) laboratory scientists and (**C**) BashTheBug volunteers . NR includes both plates that could not be read due to issues with the control wells and problems with individual drugs such as skip wells. (**D**) A heatmap showing how for each set of images assessed by the laboratory scientist has having a specific dilution as the MIC, the classifications made by BashTheBug volunteers varied considerably. It is normalised so that each row sums to 100 % and only cells with $> 0.1$ % are labelled.

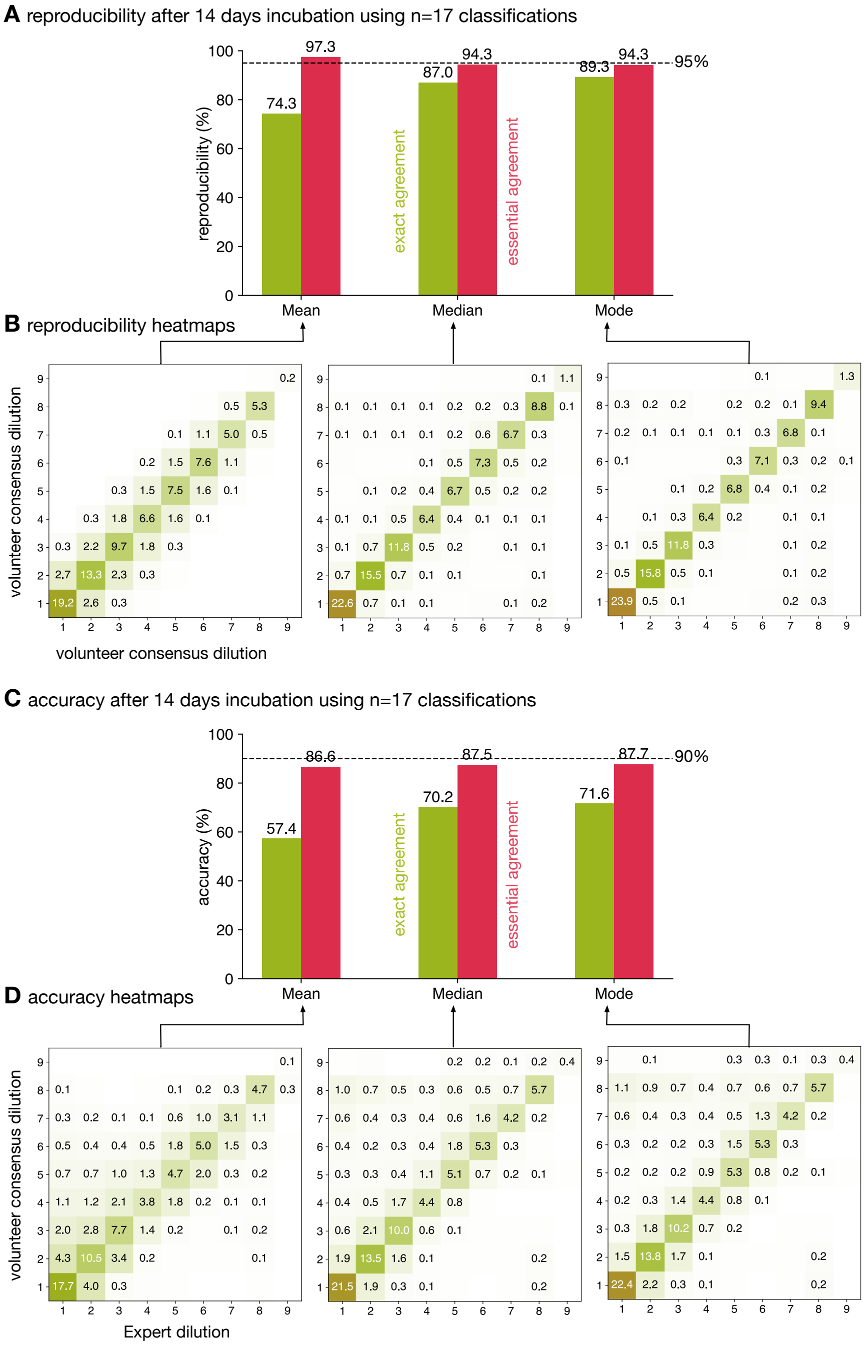

Figure S10: Taking the mean of 17 classifications is $\geq 95\%$ reproducible whilst none of the methods reach have an essential agreement for accuracy of $\geq 90\%$ when using the Expert dataset. (**A**) Only calculating the mean of 17 classifications achieves an essential agreement $\geq 95\%$ for reproducibility[1], followed by the median and then the mode. There is no specified threshold for exact agreement; the trend is reversed with the mode performing best, followed by the median and then the mean. (**B**) Heatmaps of the consensus formed via the mean, median or mode after 14 days incubation. Each consensus dilution is a different selection, with replacement, of the original classifications. Drug images from the larger Expert dataset are included. (**C**) The essential agreement between a consensus dilution formed from 17 classifications using the median or mode and the consensus Expert dilution is $\geq 90\%$, which is the required threshold[1]. (**D**) The heatmaps clearly show how the volunteer consensus dilution is likely to be the same or greater than the Expert consensus.

| Reading day | $n$ | Prop. with MIC (%) | Exact Agreement (%) Mean | Exact Agreement (%) Median | Exact Agreement (%) Mode | Essential Agreement (%) Mean | Essential Agreement (%) Median | Essential Agreement (%) Mode |
|---|---|---|---|---|---|---|---|---|
| | 1 | $62.6 \pm 0.1$ | $68.6 \pm 0.1$ | $68.6 \pm 0.1$ | $68.6 \pm 0.1$ | $82.6 \pm 0.1$ | $82.6 \pm 0.1$ | $82.6 \pm 0.1$ |
| | 3 | $69.3 \pm 0.1$ | $61.7 \pm 0.1$ | $71.6 \pm 0.1$ | $75.2 \pm 0.1$ | $84.3 \pm 0.1$ | $86.0 \pm 0.1$ | $86.9 \pm 0.1$ |
| | 5 | $71.8 \pm 0.1$ | $63.3 \pm 0.1$ | $76.9 \pm 0.1$ | $79.4 \pm 0.1$ | $89.5 \pm 0.1$ | $89.5 \pm 0.1$ | $89.1 \pm 0.1$ |
| | 7 | $73.3 \pm 0.1$ | $65.5 \pm 0.1$ | $79.2 \pm 0.1$ | $82.1 \pm 0.1$ | $92.5 \pm 0.1$ | $91.0 \pm 0.1$ | $90.8 \pm 0.1$ |
| 7 | 9 | $74.4 \pm 0.1$ | $67.7 \pm 0.1$ | $81.5 \pm 0.1$ | $84.1 \pm 0.1$ | $94.0 \pm 0.1$ | $92.2 \pm 0.1$ | $91.7 \pm 0.1$ |
| | 11 | $75.2 \pm 0.1$ | $69.8 \pm 0.1$ | $82.8 \pm 0.1$ | $85.4 \pm 0.1$ | $95.2 \pm 0.1$ | $93.0 \pm 0.1$ | $92.6 \pm 0.1$ |
| | 13 | $75.8 \pm 0.1$ | $71.6 \pm 0.1$ | $84.2 \pm 0.1$ | $86.7 \pm 0.1$ | $96.1 \pm 0.1$ | $93.6 \pm 0.1$ | $93.2 \pm 0.1$ |
| | 15 | $76.2 \pm 0.1$ | $72.6 \pm 0.1$ | $85.1 \pm 0.1$ | $87.8 \pm 0.1$ | $96.7 \pm 0.1$ | $94.2 \pm 0.1$ | $93.7 \pm 0.1$ |
| | 17 | $76.4 \pm 0.1$ | $73.6 \pm 0.1$ | $85.4 \pm 0.1$ | $87.9 \pm 0.1$ | $97.2 \pm 0.1$ | $94.3 \pm 0.1$ | $93.8 \pm 0.1$ |
| | 1 | $75.5 \pm 0.1$ | $68.7 \pm 0.1$ | $68.7 \pm 0.1$ | $68.7 \pm 0.1$ | $82.9 \pm 0.1$ | $82.9 \pm 0.1$ | $82.9 \pm 0.1$ |
| | 3 | $82.1 \pm 0.1$ | $62.5 \pm 0.1$ | $72.1 \pm 0.1$ | $75.8 \pm 0.1$ | $84.9 \pm 0.1$ | $85.9 \pm 0.1$ | $87.4 \pm 0.1$ |
| | 5 | $85.3 \pm 0.1$ | $63.7 \pm 0.1$ | $77.3 \pm 0.1$ | $79.8 \pm 0.1$ | $89.9 \pm 0.1$ | $89.4 \pm 0.1$ | $89.6 \pm 0.1$ |
| | 7 | $86.6 \pm 0.1$ | $66.1 \pm 0.1$ | $79.4 \pm 0.1$ | $82.4 \pm 0.1$ | $92.7 \pm 0.1$ | $90.7 \pm 0.1$ | $90.8 \pm 0.1$ |
| 10 | 9 | $87.5 \pm 0.1$ | $68.2 \pm 0.1$ | $81.5 \pm 0.1$ | $84.2 \pm 0.1$ | $94.1 \pm 0.1$ | $91.8 \pm 0.1$ | $91.7 \pm 0.1$ |
| | 11 | $88.2 \pm 0.1$ | $70.1 \pm 0.1$ | $82.9 \pm 0.1$ | $85.4 \pm 0.1$ | $95.3 \pm 0.1$ | $92.6 \pm 0.1$ | $92.3 \pm 0.1$ |
| | 13 | $88.7 \pm 0.1$ | $71.8 \pm 0.1$ | $84.1 \pm 0.1$ | $86.5 \pm 0.1$ | $96.1 \pm 0.1$ | $93.2 \pm 0.1$ | $93.0 \pm 0.1$ |
| | 15 | $89.1 \pm 0.1$ | $73.1 \pm 0.1$ | $84.9 \pm 0.1$ | $87.3 \pm 0.1$ | $96.8 \pm 0.1$ | $93.6 \pm 0.1$ | $93.3 \pm 0.1$ |
| | 17 | $89.9 \pm 0.1$ | $74.0 \pm 0.1$ | $85.5 \pm 0.1$ | $88.0 \pm 0.1$ | $97.2 \pm 0.1$ | $93.9 \pm 0.1$ | $93.7 \pm 0.1$ |
| | 1 | $76.5 \pm 0.1$ | $70.4 \pm 0.1$ | $70.4 \pm 0.1$ | $70.4 \pm 0.1$ | $83.2 \pm 0.1$ | $83.2 \pm 0.1$ | $83.2 \pm 0.1$ |
| | 3 | $83.9 \pm 0.1$ | $63.9 \pm 0.1$ | $73.8 \pm 0.1$ | $77.9 \pm 0.1$ | $85.2 \pm 0.1$ | $86.4 \pm 0.1$ | $88.4 \pm 0.1$ |
| | 5 | $87.0 \pm 0.1$ | $64.5 \pm 0.1$ | $78.8 \pm 0.1$ | $81.7 \pm 0.1$ | $90.0 \pm 0.1$ | $89.7 \pm 0.1$ | $90.4 \pm 0.1$ |
| | 7 | $88.7 \pm 0.1$ | $66.5 \pm 0.1$ | $80.9 \pm 0.1$ | $84.0 \pm 0.1$ | $92.6 \pm 0.1$ | $90.9 \pm 0.1$ | $91.5 \pm 0.1$ |
| 14 | 9 | $89.8 \pm 0.1$ | $68.4 \pm 0.1$ | $82.9 \pm 0.1$ | $85.5 \pm 0.1$ | $94.2 \pm 0.1$ | $92.0 \pm 0.1$ | $92.2 \pm 0.1$ |
| | 11 | $90.3 \pm 0.1$ | $70.1 \pm 0.1$ | $84.3 \pm 0.1$ | $87.0 \pm 0.1$ | $95.3 \pm 0.1$ | $92.9 \pm 0.1$ | $93.0 \pm 0.1$ |
| | 13 | $91.0 \pm 0.1$ | $71.6 \pm 0.1$ | $85.1 \pm 0.1$ | $87.7 \pm 0.1$ | $96.1 \pm 0.1$ | $93.4 \pm 0.1$ | $93.4 \pm 0.1$ |
| | 15 | $91.2 \pm 0.1$ | $72.9 \pm 0.1$ | $86.0 \pm 0.1$ | $88.4 \pm 0.1$ | $96.8 \pm 0.1$ | $93.8 \pm 0.1$ | $93.8 \pm 0.1$ |
| | 17 | $92.0 \pm 0.1$ | $74.3 \pm 0.1$ | $86.9 \pm 0.1$ | $89.2 \pm 0.1$ | $97.2 \pm 0.1$ | $94.2 \pm 0.1$ | $94.1 \pm 0.1$ |
| | 1 | $79.5 \pm 0.1$ | $74.1 \pm 0.1$ | $74.1 \pm 0.1$ | $74.1 \pm 0.1$ | $85.2 \pm 0.1$ | $85.2 \pm 0.1$ | $85.2 \pm 0.1$ |
| | 3 | $86.3 \pm 0.1$ | $68.0 \pm 0.1$ | $77.4 \pm 0.1$ | $81.4 \pm 0.1$ | $87.0 \pm 0.1$ | $88.0 \pm 0.1$ | $90.1 \pm 0.1$ |
| | 5 | $89.0 \pm 0.1$ | $68.6 \pm 0.1$ | $82.3 \pm 0.1$ | $85.0 \pm 0.1$ | $91.8 \pm 0.1$ | $91.1 \pm 0.1$ | $91.9 \pm 0.1$ |
| | 7 | $90.5 \pm 0.1$ | $70.7 \pm 0.1$ | $84.2 \pm 0.1$ | $87.0 \pm 0.1$ | $93.9 \pm 0.1$ | $92.1 \pm 0.1$ | $92.8 \pm 0.1$ |
| 21 | 9 | $91.5 \pm 0.1$ | $72.4 \pm 0.1$ | $85.7 \pm 0.1$ | $88.1 \pm 0.1$ | $95.1 \pm 0.1$ | $93.1 \pm 0.1$ | $93.4 \pm 0.1$ |
| | 11 | $92.0 \pm 0.1$ | $73.7 \pm 0.1$ | $87.0 \pm 0.1$ | $89.3 \pm 0.1$ | $96.2 \pm 0.1$ | $93.7 \pm 0.1$ | $94.0 \pm 0.1$ |
| | 13 | $92.4 \pm 0.1$ | $75.0 \pm 0.1$ | $87.8 \pm 0.1$ | $90.1 \pm 0.1$ | $96.7 \pm 0.1$ | $94.2 \pm 0.1$ | $94.4 \pm 0.1$ |
| | 15 | $92.7 \pm 0.1$ | $76.1 \pm 0.1$ | $88.4 \pm 0.1$ | $90.5 \pm 0.1$ | $97.4 \pm 0.1$ | $94.6 \pm 0.1$ | $94.6 \pm 0.1$ |
| | 17 | $93.3 \pm 0.1$ | $77.3 \pm 0.1$ | $88.9 \pm 0.1$ | $91.2 \pm 0.1$ | $97.7 \pm 0.1$ | $94.9 \pm 0.1$ | $95.0 \pm 0.1$ |

Table S5: The effect on reproducibility of varying the number of days incubated, the number of classifications, $n$, and the consensus method.

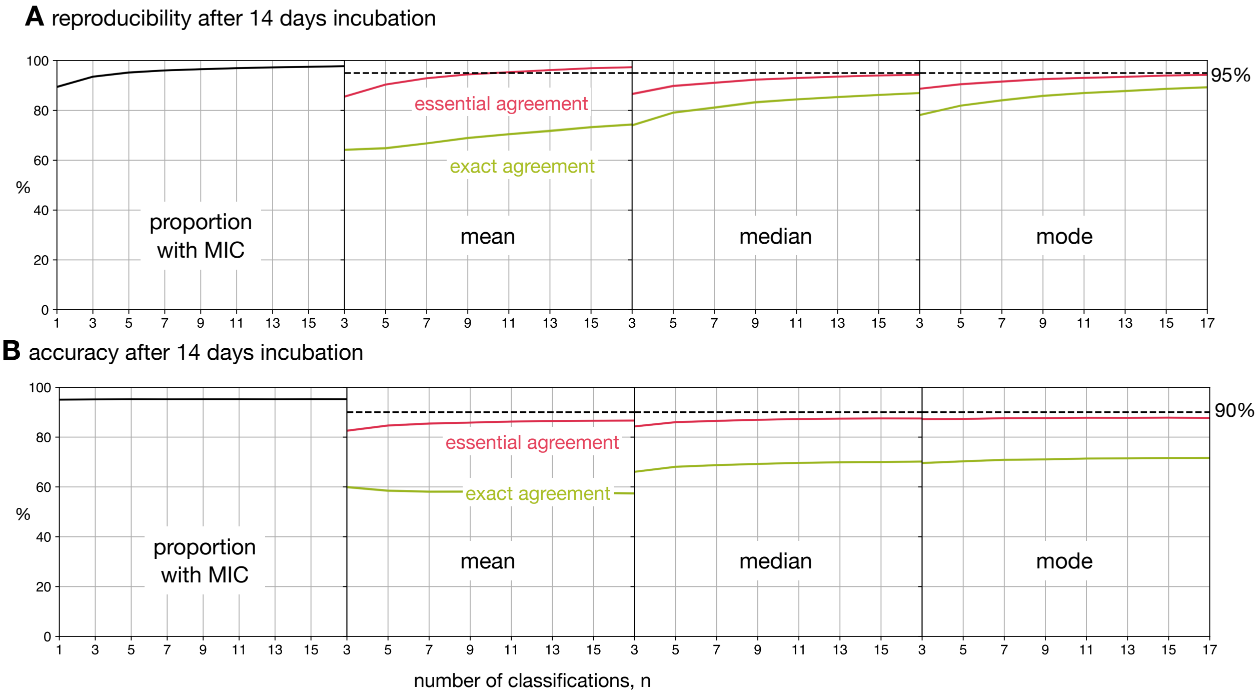

Figure S11: Reducing the number of classifications, $n$, used to build the consensus dilution decreases the reproducibility and accuracy of the consensus measurement. (**A**) The consensus dilution becomes less reproducible as the number of classifications is reduced, as measured by both the exact and essential agreements. (**B**) Likewise, the consensus dilution becomes less accurate as the number of classifications is decreased, however the highest level of exact agreement using the mean is obtained when $n = 3$ and the mode, and to a lesser extent the median, are relatively insensitive to the number of classifications. These data are all with respect to the Expert dataset.

| Reading day | $n$ | Prop. with MIC (%) | Exact Agreement (%) Mean | Median | Mode | Essential Agreement (%) Mean | Median | Mode |
|---|---|---|---|---|---|---|---|---|
| 7 | 3 | $78.0 \pm 0.1$ | $67.6 \pm 0.2$ | $69.1 \pm 0.2$ | $74.0 \pm 0.2$ | $84.0 \pm 0.1$ | $85.8 \pm 0.1$ | $89.4 \pm 0.1$ |
| | 5 | $78.5 \pm 0.1$ | $67.3 \pm 0.1$ | $70.1 \pm 0.1$ | $73.5 \pm 0.1$ | $86.5 \pm 0.1$ | $87.3 \pm 0.1$ | $89.2 \pm 0.1$ |
| | 7 | $78.9 \pm 0.1$ | $67.9 \pm 0.2$ | $70.3 \pm 0.1$ | $73.7 \pm 0.1$ | $87.4 \pm 0.1$ | $87.6 \pm 0.1$ | $89.2 \pm 0.1$ |
| | 9 | $78.9 \pm 0.1$ | $68.7 \pm 0.1$ | $70.6 \pm 0.1$ | $73.7 \pm 0.1$ | $87.6 \pm 0.1$ | $88.0 \pm 0.1$ | $89.2 \pm 0.1$ |
| | 11 | $79.2 \pm 0.1$ | $69.1 \pm 0.2$ | $70.8 \pm 0.1$ | $73.8 \pm 0.1$ | $88.4 \pm 0.1$ | $88.3 \pm 0.1$ | $89.2 \pm 0.1$ |
| | 13 | $79.2 \pm 0.1$ | $69.0 \pm 0.1$ | $70.7 \pm 0.1$ | $73.6 \pm 0.1$ | $88.6 \pm 0.1$ | $88.3 \pm 0.1$ | $89.1 \pm 0.1$ |
| | 15 | $79.2 \pm 0.1$ | $68.8 \pm 0.2$ | $70.6 \pm 0.1$ | $73.5 \pm 0.1$ | $88.7 \pm 0.1$ | $88.3 \pm 0.1$ | $89.1 \pm 0.1$ |
| | 17 | $79.4 \pm 0.1$ | $68.9 \pm 0.1$ | $71.6 \pm 0.1$ | $74.5 \pm 0.1$ | $88.3 \pm 0.1$ | $88.4 \pm 0.1$ | $89.1 \pm 0.1$ |
| 10 | 3 | $91.7 \pm 0.1$ | $70.3 \pm 0.1$ | $73.4 \pm 0.1$ | $78.5 \pm 0.1$ | $84.0 \pm 0.1$ | $86.7 \pm 0.1$ | $90.4 \pm 0.1$ |
| | 5 | $92.0 \pm 0.1$ | $69.3 \pm 0.1$ | $75.0 \pm 0.1$ | $78.4 \pm 0.1$ | $86.3 \pm 0.1$ | $88.4 \pm 0.1$ | $90.4 \pm 0.1$ |
| | 7 | $92.0 \pm 0.1$ | $69.6 \pm 0.1$ | $75.4 \pm 0.1$ | $78.7 \pm 0.1$ | $87.3 \pm 0.1$ | $88.9 \pm 0.1$ | $90.5 \pm 0.1$ |
| | 9 | $92.1 \pm 0.1$ | $70.1 \pm 0.1$ | $75.5 \pm 0.1$ | $78.7 \pm 0.1$ | $87.4 \pm 0.1$ | $89.2 \pm 0.1$ | $90.5 \pm 0.1$ |
| | 11 | $92.2 \pm 0.1$ | $70.1 \pm 0.1$ | $75.8 \pm 0.1$ | $78.9 \pm 0.1$ | $87.9 \pm 0.1$ | $89.5 \pm 0.1$ | $90.6 \pm 0.1$ |
| | 13 | $92.2 \pm 0.1$ | $70.0 \pm 0.1$ | $75.9 \pm 0.1$ | $78.7 \pm 0.1$ | $88.3 \pm 0.1$ | $89.7 \pm 0.1$ | $90.5 \pm 0.1$ |
| | 15 | $92.1 \pm 0.1$ | $69.8 \pm 0.1$ | $75.9 \pm 0.1$ | $78.9 \pm 0.1$ | $88.4 \pm 0.1$ | $89.8 \pm 0.1$ | $90.6 \pm 0.1$ |
| | 17 | $92.5 \pm 0.1$ | $69.6 \pm 0.1$ | $75.9 \pm 0.1$ | $78.7 \pm 0.1$ | $88.0 \pm 0.1$ | $89.5 \pm 0.1$ | $90.3 \pm 0.1$ |
| 14 | 3 | $97.1 \pm 0.1$ | $69.9 \pm 0.1$ | $74.5 \pm 0.1$ | $80.0 \pm 0.1$ | $83.6 \pm 0.1$ | $86.7 \pm 0.1$ | $90.6 \pm 0.1$ |
| | 5 | $97.2 \pm 0.1$ | $68.4 \pm 0.1$ | $76.4 \pm 0.1$ | $80.1 \pm 0.1$ | $86.3 \pm 0.1$ | $88.6 \pm 0.1$ | $90.7 \pm 0.1$ |
| | 7 | $97.3 \pm 0.1$ | $68.1 \pm 0.1$ | $76.9 \pm 0.1$ | $80.5 \pm 0.1$ | $87.0 \pm 0.1$ | $89.0 \pm 0.1$ | $90.9 \pm 0.1$ |
| | 9 | $97.3 \pm 0.1$ | $68.6 \pm 0.1$ | $77.2 \pm 0.1$ | $80.5 \pm 0.1$ | $87.5 \pm 0.1$ | $89.6 \pm 0.1$ | $90.9 \pm 0.1$ |
| | 11 | $97.3 \pm 0.1$ | $68.9 \pm 0.1$ | $77.7 \pm 0.1$ | $80.9 \pm 0.1$ | $87.9 \pm 0.1$ | $89.9 \pm 0.1$ | $91.1 \pm 0.1$ |
| | 13 | $97.4 \pm 0.1$ | $68.7 \pm 0.1$ | $77.9 \pm 0.1$ | $80.9 \pm 0.1$ | $88.3 \pm 0.1$ | $90.1 \pm 0.1$ | $91.1 \pm 0.1$ |
| | 15 | $97.4 \pm 0.1$ | $68.5 \pm 0.1$ | $78.0 \pm 0.1$ | $80.9 \pm 0.1$ | $88.4 \pm 0.1$ | $90.2 \pm 0.1$ | $91.1 \pm 0.1$ |
| | 17 | $97.4 \pm 0.1$ | $68.4 \pm 0.1$ | $78.1 \pm 0.1$ | $80.9 \pm 0.1$ | $88.5 \pm 0.1$ | $90.2 \pm 0.1$ | $91.0 \pm 0.1$ |
| 21 | 3 | $98.9 \pm 0.1$ | $74.3 \pm 0.1$ | $79.3 \pm 0.1$ | $84.0 \pm 0.1$ | $86.3 \pm 0.1$ | $89.0 \pm 0.1$ | $92.3 \pm 0.1$ |
| | 5 | $99.0 \pm 0.1$ | $73.1 \pm 0.1$ | $81.3 \pm 0.1$ | $84.5 \pm 0.1$ | $88.8 \pm 0.1$ | $90.7 \pm 0.1$ | $92.5 \pm 0.1$ |
| | 7 | $99.0 \pm 0.1$ | $73.4 \pm 0.1$ | $81.9 \pm 0.1$ | $85.1 \pm 0.1$ | $89.6 \pm 0.1$ | $91.4 \pm 0.1$ | $92.9 \pm 0.1$ |
| | 9 | $99.0 \pm 0.1$ | $73.7 \pm 0.1$ | $82.4 \pm 0.1$ | $85.2 \pm 0.1$ | $89.8 \pm 0.1$ | $91.7 \pm 0.1$ | $92.9 \pm 0.1$ |
| | 11 | $99.0 \pm 0.1$ | $73.5 \pm 0.1$ | $82.7 \pm 0.1$ | $85.5 \pm 0.1$ | $90.2 \pm 0.1$ | $92.1 \pm 0.1$ | $93.1 \pm 0.1$ |
| | 13 | $99.0 \pm 0.1$ | $73.8 \pm 0.1$ | $83.0 \pm 0.1$ | $85.6 \pm 0.1$ | $90.5 \pm 0.1$ | $92.4 \pm 0.1$ | $93.2 \pm 0.1$ |
| | 15 | $99.0 \pm 0.1$ | $73.5 \pm 0.1$ | $83.2 \pm 0.1$ | $85.6 \pm 0.1$ | $90.6 \pm 0.1$ | $92.5 \pm 0.1$ | $93.2 \pm 0.1$ |
| | 17 | $99.0 \pm 0.1$ | $73.3 \pm 0.1$ | $83.2 \pm 0.1$ | $85.6 \pm 0.1$ | $90.6 \pm 0.1$ | $92.6 \pm 0.1$ | $93.3 \pm 0.1$ |

Table S6: The effect on accuracy of varying the number of days incubated, the number of classifications, $n$, and the consensus method.

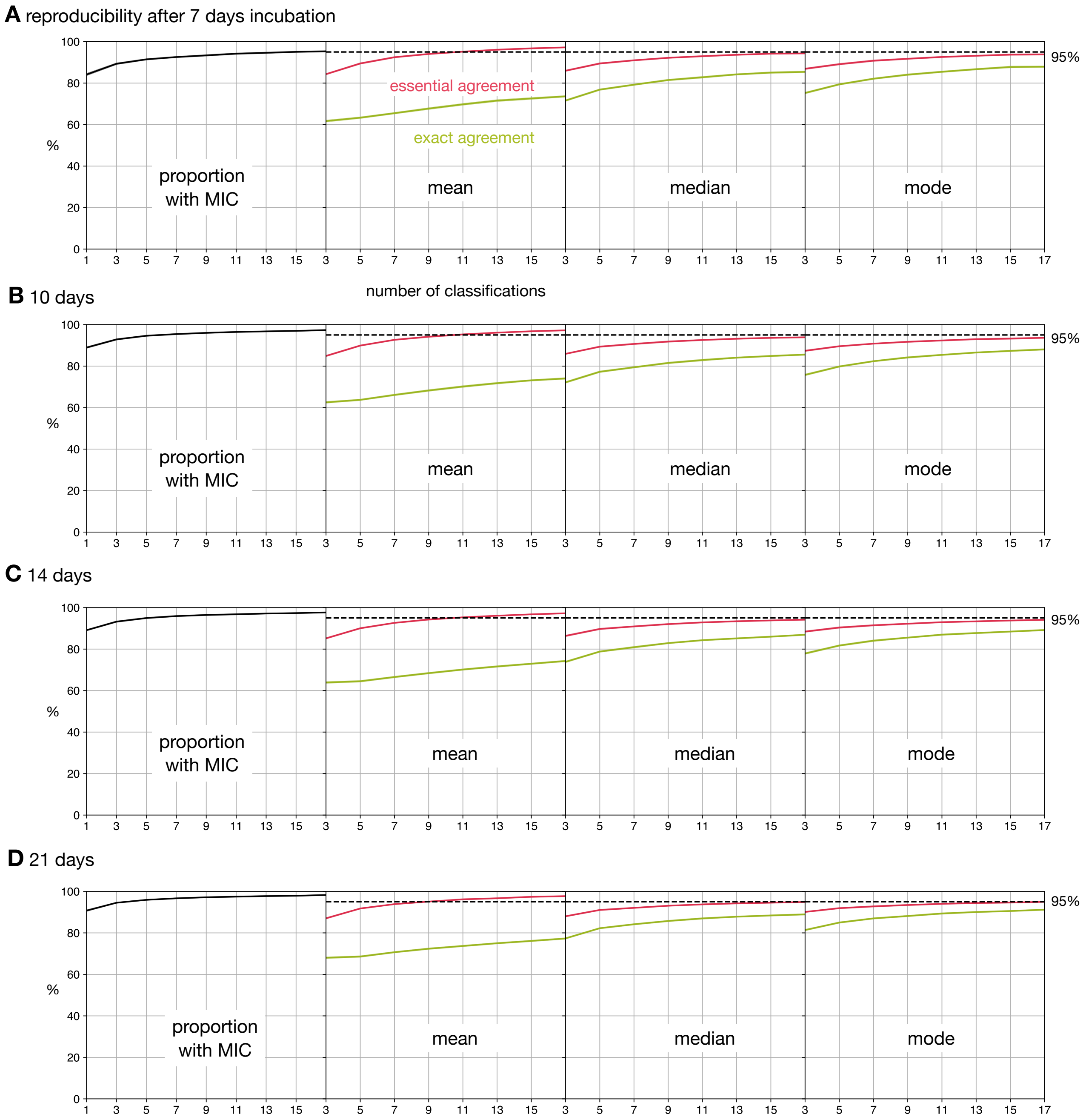

Figure S12: Altering the number of days incubation does not markedly affect the observed trends in reproducibility. Shown are results for the Expert+AMyGDA dataset after (**A**) 7, (**B**) 10, (**C**) 14 and (**D**) 21 days of incubation. A previous study[2] showed that optimal results were achieved after 14 days incubation.

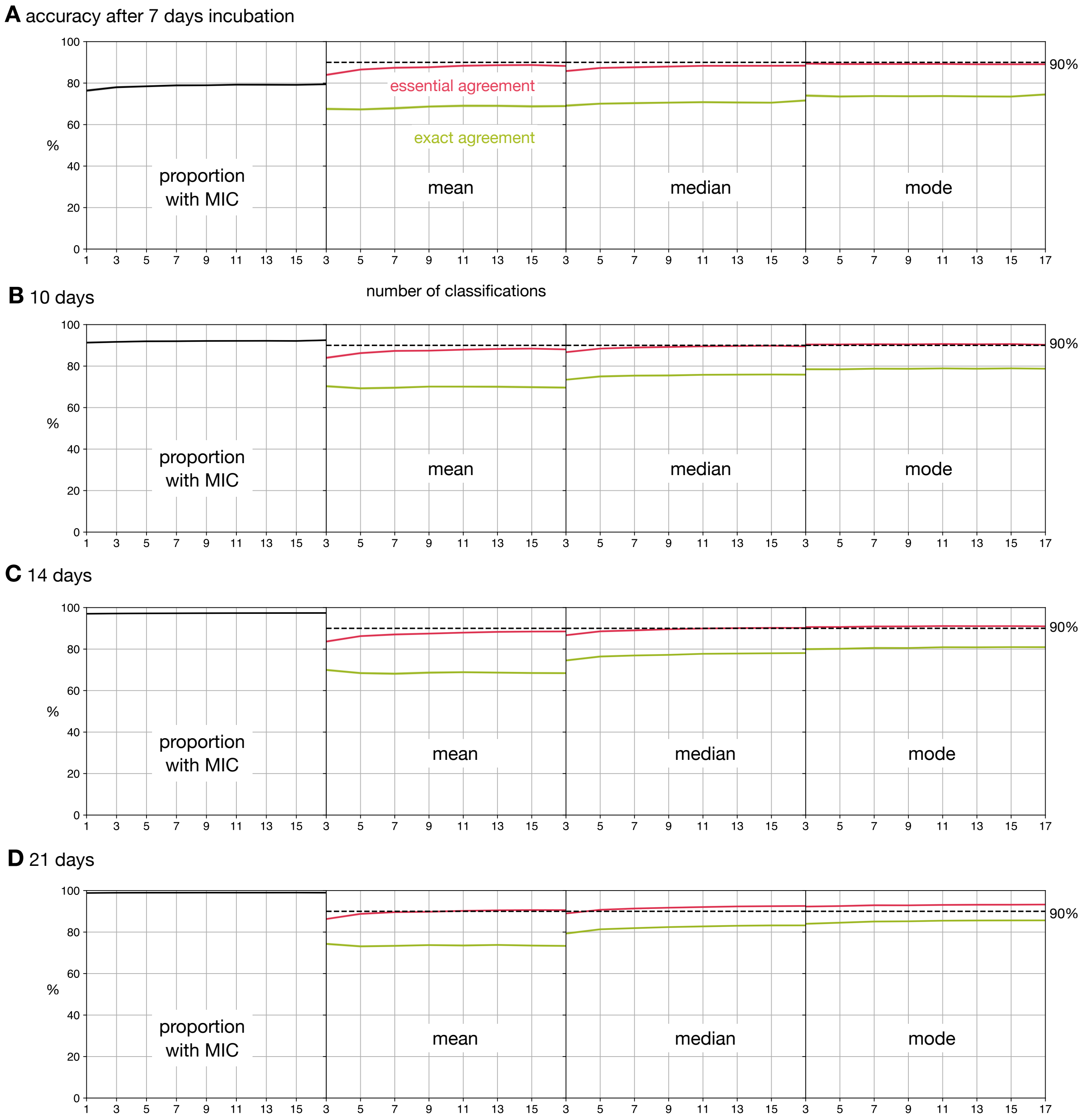

Figure S13: Altering the number of days incubation does not markedly affect the observed trends in accuracy. Shown are results for the Expert+AMyGDA dataset after (**A**) 7, (**B**) 10, (**C**) 14 and (**D**) 21 days of incubation. A previous study[2] showed that optimal results were achieved after 14 days incubation.

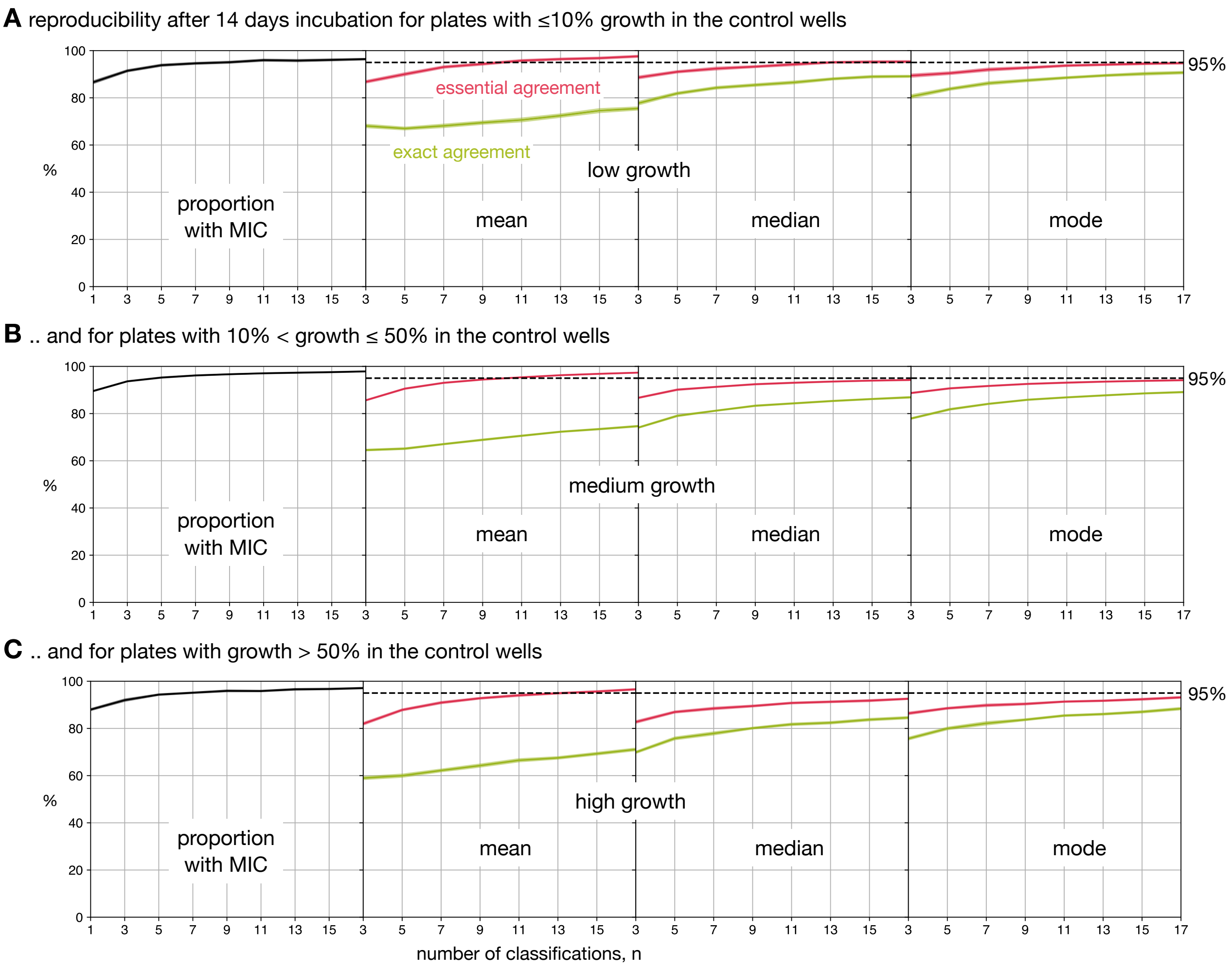

Figure S14: Segmenting the drug images by the mean amount of growth in the positive control wells (Fig. S7) does not markedly affect the reproducibility of the three consensus methods. The plates are split into those with (**A**) low ($\leq 10$ %) growth, (**B**) medium ($10 <$ growth $\leq 50\%$) growth and (**C**) high ($> 50\%$) growth. The drug images from the Expert+AMyGDA dataset were used and the proportion with MIC is the proportion of consensus readings that are a definite numerical minimum inhibitory concentration.

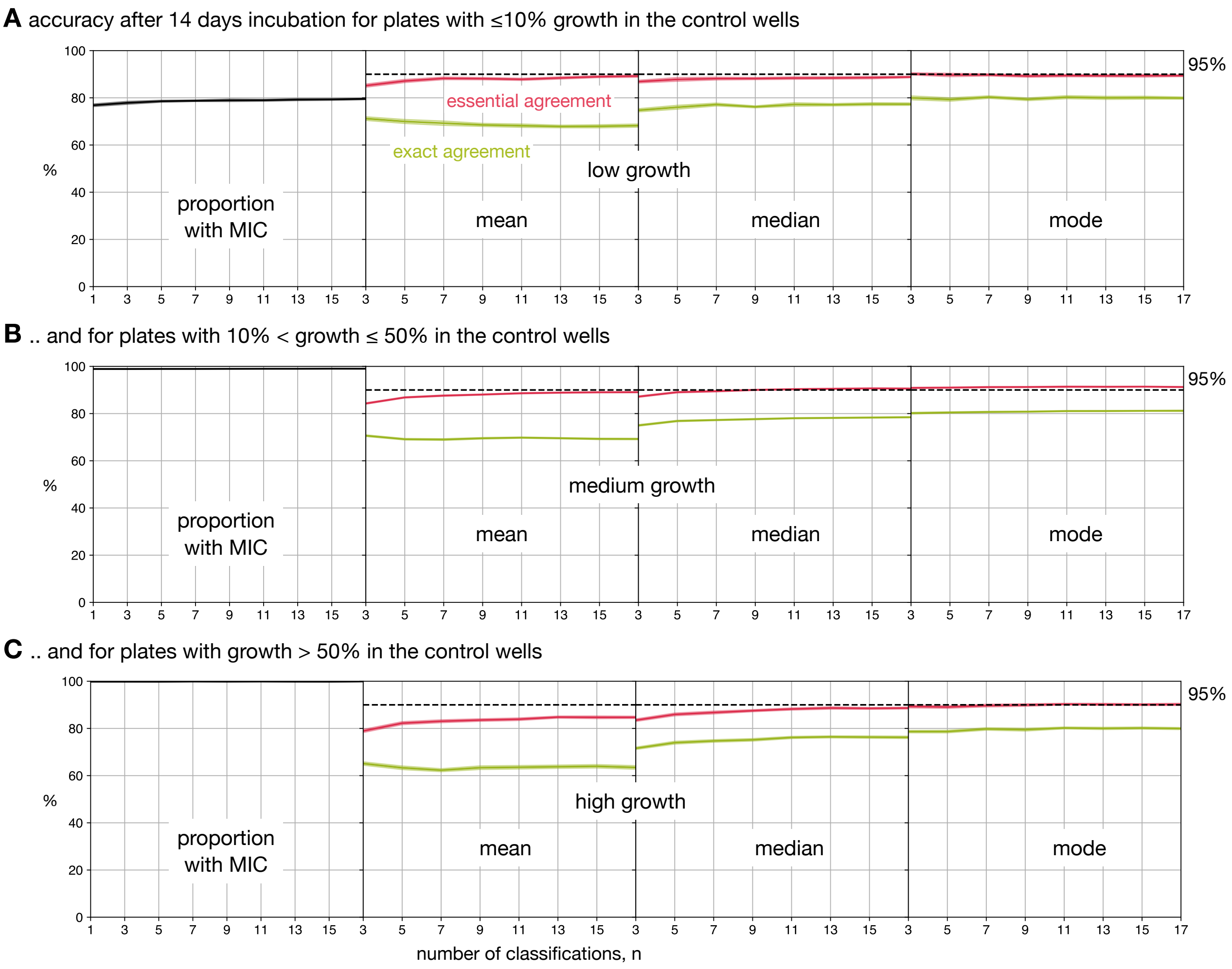

Figure S15: Segmenting the drug images by the mean amount of growth in the positive control wells (Fig. S7) does not markedly affect the accuracy of the three consensus methods. The plates are split into those with (**A**) low ($\leq 10$ %) growth, (**B**) medium ($10 <$ growth $\leq 50\%$) growth and (**C**) high ($> 50\%$) growth. The drug images from the Expert+AMyGDA dataset were used and the proportion with MIC is the proportion of consensus readings that are a definite numerical minimum inhibitory concentration.

| Reading day | n | Average n | Prop. with MIC (%) | Exact Agreement (%) Mean | Exact Agreement (%) Median | Exact Agreement (%) Mode | Essential Agreement (%) Mean | Essential Agreement (%) Median | Essential Agreement (%) Mode |
|---|---|---|---|---|---|---|---|---|---|
| 14 | 3 | 3.0 | 95.1 | 81.4 | 79.1 | 81.7 | 92.6 | 89.3 | 90.4 |
| | 5 | 3.9 | 96.6 | 79.0 | 82.7 | 84.4 | 93.1 | 91.4 | 91.6 |
| | 7 | 4.7 | 97.1 | 80.0 | 83.6 | 85.7 | 94.4 | 92.1 | 92.1 |
| | 9 | 5.6 | 97.4 | 80.8 | 84.6 | 86.6 | 95.3 | 92.8 | 92.7 |
| | 11 | 6.5 | 97.6 | 82.0 | 85.5 | 87.3 | 96.1 | 93.3 | 93.1 |
| | 13 | 7.3 | 97.8 | 82.8 | 86.1 | 88.0 | 96.6 | 93.7 | 93.4 |
| | 15 | 8.2 | 98.0 | 83.3 | 86.6 | 88.5 | 96.9 | 93.9 | 93.7 |
| | 17 | 8.8 | 98.2 | 84.5 | 87.6 | 89.4 | 97.3 | 94.4 | 94.1 |

Table S7: The effect on reproducibility of dynamically retiring images if the first three classifications are identical and continuing the remainder until they have accrued *n* classifications.

| Reading day | n | Average n | Prop. with MIC (%) | Exact Agreement (%) Mean | Exact Agreement (%) Median | Exact Agreement (%) Mode | Essential Agreement (%) Mean | Essential Agreement (%) Median | Essential Agreement (%) Mode |
|---|---|---|---|---|---|---|---|---|---|
| 14 | 3 | 3.0 | 97.2 | 69.9 | 74.5 | 80.0 | 83.6 | 86.7 | 90.6 |
| | 5 | 3.8 | 96.9 | 74.4 | 77.7 | 80.2 | 88.1 | 89.2 | 90.6 |
| | 7 | 4.6 | 97.0 | 74.1 | 77.8 | 80.4 | 88.5 | 89.5 | 90.7 |
| | 9 | 5.4 | 97.0 | 74.2 | 77.9 | 80.4 | 88.6 | 89.8 | 90.7 |
| | 11 | 6.2 | 97.0 | 74.4 | 78.2 | 80.6 | 88.9 | 90.0 | 90.8 |
| | 13 | 7.0 | 97.1 | 74.4 | 78.3 | 80.5 | 89.1 | 90.1 | 90.8 |
| | 15 | 7.8 | 97.1 | 74.2 | 78.3 | 80.6 | 89.1 | 90.2 | 90.8 |
| | 17 | 8.4 | 97.1 | 74.8 | 78.8 | 80.9 | 89.4 | 90.3 | 90.9 |

Table S8: The effect on accuracy of dynamically retiring images if the first three classifications are identical and continuing the remainder until they have accrued *n* classifications.

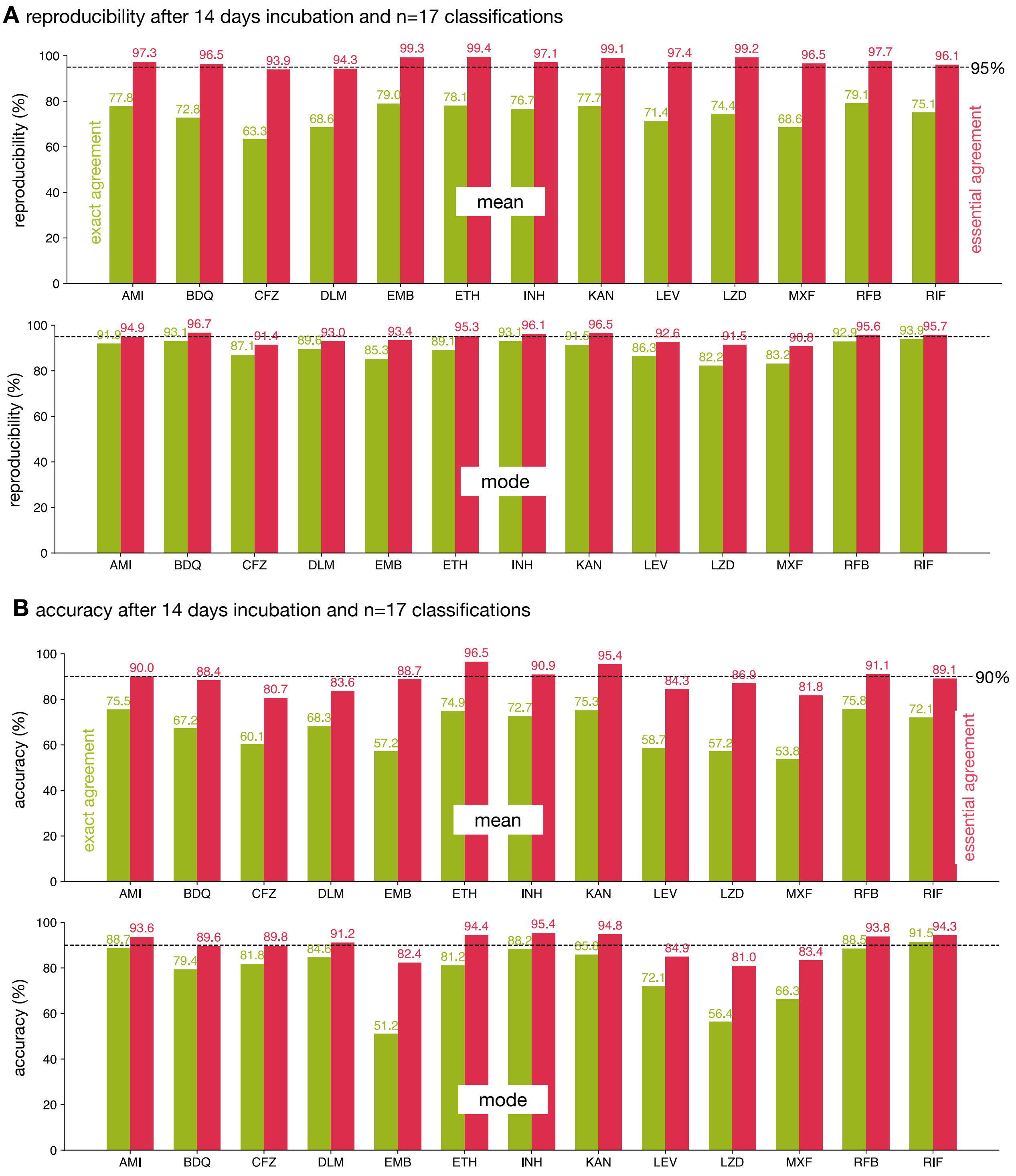

Figure S16: The reproducibility and accuracy after 14 days incubation of the 13 antibiotics on the UKMYC5 plate. A total of 17 classifications were used for each measurement and either the mean or mode was used to obtain a consensus reading of the (**A**) reproducibility and (**B**) accuracy. The essential agreement is drawn in red and the required thresholds are 95% and 90% for the reproducibility and accuracy, respectively[1]. The exact agreement is drawn in green and no threshold is defined. The drug abbreviations are defined in Fig. S2. The dataset used was Expert+AMyGDA.

| Dilution | Agreement |
|---:|---:|
| NR | 20.0 ± 0.1 % |
| 1 | 83.9 ± 0.1 % |
| 2 | 72.0 ± 0.1 % |
| 3 | 66.6 ± 0.1 % |
| 4 | 48.2 ± 0.1 % |
| 5 | 53.4 ± 0.1 % |
| 6 | 52.6 ± 0.1 % |
| 7 | 42.8 ± 0.1 % |
| 8 | 46.7 ± 0.1 % |
| 9 | 21.1 ± 0.2 % |

Table S9: The Expert and BashTheBug MICs are more likely to concur at smaller dilutions. The BashTheBug consensus measurement was built by taking the median of 17 classifications and rounding up if a non-integer was returned.

### MEMBERS OF CRyPTIC CONSORTIUM

Ivan Barilar[29], Simone Battaglia[1], Emanuele Borroni[1], Angela Pires Brandao[2,3], Alice Brankin[4], Andrea Maurizio Cabibbe[1], Joshua Carter[5], Daniela Maria Cirillo[1], Pauline Claxton[6], David A Clifton[4], Ted Cohen[7], Jorge Coronel[8], Derrick W Crook[4], Viola Dreyer[29], Sarah G Earle[4], Vincent Escuyer[9], Lucilaine Ferrazoli[3], Philip W Fowler[4], George Fu Gao[10], Jennifer Gardy[11], Saheer Gharbia[12], Kelen Teixeira Ghisi[3], Arash Ghodousi[1,13], Ana Luíza Gibertoni Cruz[4], Louis Grandjean[33], Clara Grazian[14], Ramona Groenheit[44], Jennifer L Guthrie[15,16], Wencong He[10], Harald Hoffmann[17,18], Sarah J Hoosdally[4], Martin Hunt[4,19], Zamin Iqbal[19], Nazir Ahmed Ismail[20], Lisa Jarrett[21], Lavania Joseph[20], Ruwen Jou[22], Priti Kambli[23], Rukhsar Khot[23], Jeff Knaggs[4,19], Anastasia Koch[24], Donna Kohlerschmidt[9], Samaneh Kouchaki[4,25], Alexander S Lachapelle[4], Ajit Lalvani[26], Simon Grandjean Lapierre[27], Ian F Laurenson[6], Brice Letcher[19], Wan-Hsuan Lin[22], Chunfa Liu[10], Dongxin Liu[10], Kerri M Malone[19], Ayan Mandal[28], Mikael Mansjö[44], Daniela Matias[21], Graeme Meintjes[24], Flávia de Freitas Mendes[3], Matthias Merker[29], Marina Mihalic[18], James Millard[30], Paolo Miotto[1], Nerges Mistry[28], David Moore[8,31], Kimberlee A Musser[9], Dumisani Ngcamu[20], Hoang Ngoc Nhung[32], Stefan Niemann[29,48], Kayzad Soli Nilgiriwala[28], Camus Nimmo[33], Nana Okozi[20], Rosangela Siqueira Oliveira[3], Shaheed Vally Omar[20], Nicholas Paton[34], Timothy EA Peto[4], Juliana Maira Watanabe Pinhata[3], Sara Plesnik[18], Zully M Puyen[35], Marie Sylvianne Rabodoarivelo[36], Niaina Rakotosamimanana[36], Paola MV Rancoita[13], Priti Rathod[21], Esther Robinson[21], Gillian Rodger[4], Camilla Rodrigues[23], Timothy C Rodwell[37,38], Aysha Roohi[4], David Santos-Lazaro[35], Sanchi Shah[28], Thomas Andreas Kohl[29], Grace Smith[12,21], Walter Solano[8], Andrea Spitaleri[1,13], Philip Supply[39], Utkarsha Surve[23], Sabira Tahseen[40], Nguyen Thuy Thuong Thuong[32], Guy Thwaites[4,32], Katharina Todt[18], Alberto Trovato[1], Christian Utpatel[29], Annelies Van Rie[41], Srinivasan Vijay[42], Timothy M Walker[4,32], A Sarah Walker[4], Robin Warren[43], Jim Werngren[44], Maria Wijkander[44], Robert J Wilkinson[26,45,46], Daniel J Wilson[4], Penelope Wintringer[19], Yu-Xin Xiao[22], Yang Yang[4], Zhao Yanlin[10], Shen-Yuan Yao[20], Baoli Zhu[47].

#### Affiliations

1. IRCCS San Raffaele Scientific Institute, Milan, Italy
2. Oswaldo Cruz Foundation, Rio de Janeiro, Brazil
3. Institute Adolfo Lutz, São Paulo, Brazil
4. University of Oxford, Oxford, UK
5. Stanford University School of Medicine, Stanford, USA
6. Scottish Mycobacteria Reference Laboratory, Edinburgh, UK
7. Yale School of Public Health, Yale, USA
8. Universidad Peruana Cayetano Heredia, Lima, Perú
9. Wadsworth Center, New York State Department of Health, Albany, USA
10. Chinese Center for Disease Control and Prevention, Beijing, China

11. Bill & Melinda Gates Foundation, Seattle, USA
12. UK Health Security Agency, London, UK
13. Vita-Salute San Raffaele University, Milan, Italy
14. University of New South Wales, Sydney, Australia
15. The University of British Columbia, Vancouver, Canada
16. Public Health Ontario, Toronto, Canada
17. SYNLAB Gauting, Munich, Germany
18. Institute of Microbiology and Laboratory Medicine, IMLred, WHO-SRL Gauting, Germany
19. EMBL-EBI, Hinxton, UK
20. National Institute for Communicable Diseases, Johannesburg, South Africa
21. Public Health England, Birmingham, UK
22. Taiwan Centers for Disease Control, Taipei, Taiwan
23. Hinduja Hospital, Mumbai, India
24. University of Cape Town, Cape Town, South Africa
25. University of Surrey, Guildford, UK
26. Imperial College, London, UK
27. Université de Montréal, Canada
28. The Foundation for Medical Research, Mumbai, India
29. Research Center Borstel, Borstel, Germany
30. Africa Health Research Institute, Durban, South Africa
31. London School of Hygiene and Tropical Medicine, London, UK
32. Oxford University Clinical Research Unit, Ho Chi Minh City, Viet Nam
33. University College London, London, UK
34. National University of Singapore, Singapore
35. Instituto Nacional de Salud, Lima, Perú
36. Institut Pasteur de Madagascar, Antananarivo, Madagascar
37. FIND, Geneva, Switzerland

38. University of California, San Diego, USA

39. Institut Pasteur de Lille, Lille, France

40. National TB Reference Laboratory, National TB Control Program, Islamabad, Pakistan

41. University of Antwerp, Antwerp, Belgium

42. University of Edinburgh, Edinburgh, UK

43. Stellenbosch University, Cape Town, South Africa

44. Public Health Agency of Sweden, Solna, Sweden

45. Wellcome Centre for Infectious Diseases Research in Africa, Cape Town, South Africa

46. Francis Crick Institute, London, UK

47. Institute of Microbiology, Chinese Academy of Sciences, Beijing, China

48. German Center for Infection Research (DZIF), Hamburg-Lübeck-Borstel-Riems, Germany

### References

1. International Organization for Standardization (2007) ISO 20776-2: Clinical laboratory testing and in vitro diagnostic test systems. Technical report, International Standards Organization.

2. Rancoita PMV, Cugnata F, Gibertoni Cruz AL, Borroni E, Hoosdally SJ, Walker TM, Grazian C, Davies TJ, Peto TEA, Crook DW, Fowler PW, Cirillo DM, Crook DW, Peto TEA, Walker AS, Hoosdally SJ, Gibertoni Cruz AL, Grazian C, Walker TM, Fowler PW, Wilson D, Clifton D, Iqbal Z, Hunt M, Smith EG, Rathod P, Jarrett L, Matias D, Cirillo DM, Borroni E, Battaglia S, Chiacchiaretta M, De Filippo M, Cabibbe A, Tahseen S, Mistry N, Nilgiriwala K, Chitalia V, Ganesan N, Papewar A, Rodrigues C, Kambli P, Surve U, Khot R, Niemann S, Kohl T, Merker M, Hoffmann H, Lehmann S, Plesnik S, Ismail N, Omar SV, Joseph L, Marubini E, Thwaites G, Thuy Thuong TN, Ngoc NH, Srinivasan V, Moore D, Coronel J, Solano W, He G, Zhu B, Zhou Y, Ma A, Yu P, Schito M, Claxton P, Laurenson I (2018) *Antimicrobial Agents and Chemotherapy* 62:e00344–18.
